# Resting-state connectivity modifies the effects of amyloid on cognitive and physical function: evidence for network-based cognitive reserve

**DOI:** 10.1101/2024.11.07.622257

**Authors:** Paul J. Laurienti, Stephen B. Kritchevsky, Robert G. Lyday, Michael E. Miller, Samuel N. Lockhart, Melissa M. Rundle, Christina E. Hugenschmidt, Jonathan H. Burdette, Heather M. Shappell, Haiying Chen, Laura D Baker, Blake R. Neyland, Roee Holtzer

## Abstract

Cognitive and physical function are interrelated in aging co-occurring impairments in both domains can be debilitating and lead to increased risk of developing dementia. Amyloid beta (Aβ) deposition in the brain is linked to cognitive decline and is also associated with poorer physical function in older adults. However, significant inter-individual variability exists with respect to the influence of increased brain Aβ concentrations on cognitive and physical outcomes. Identifying factors that explain inter-individual variability in associations between Aβ and clinical outcomes could inform interventions designed to delay declines in both cognitive and physical function. Cognitive reserve (CR) is considered a buffer that allows for cognitive performance that is better than expected for a given level of brain injury or pathology. Although the neural mechanisms underlying CR remain unknown, there is growing evidence that resting-state brain networks may serve as a neural surrogate for CR. The currently study evaluated whether functional brain networks modified associations between brain Aβ and cognitive and physical function in community-dwelling older adults from the Brain Networks and Mobility (B-NET) study. We found that the integrity of the central executive and basal ganglia networks modified associations of Aβ with cognitive and physical performance. Associations between brain Aβ and cognitive and physical function were less pronounced when brain network integrity was high. The current study introduces novel evidence for brain networks underlying CR as a buffer against the influence of Aβ accumulation on cognitive and physical function.

**Significance Statement:** There is a growing number of medications targeting beta amyloid for the treatment of Alzheimer’s disease. The treatments effectively lower brain amyloid but do not have as robust of an effect on clinical outcomes. The current study introduces novel evidence for brain networks as a buffer against the influence of Aβ accumulation on cognitive and physical function in older adults with normal cognition. Future studies should examine if brain network integrity underlies the variability in treatment response to amyloid-lowering drugs in patients with cognitive decline.

## 1. Introduction

Among older adults, common and debilitating impairments in cognitive [1, 2] and physical [3] function result in adverse health outcomes and increased economic burden [4-7]. Furthermore, cognitive and physical function are interrelated in aging [8-10], and co-occurring impairments in both domains lead to increased risk of developing dementia [11]. Amyloid beta (Aβ) is an established biomarker of Alzheimer’s disease [12, 13] and has been linked to cognitive decline among older adults [14]. Aβ is also associated with poorer physical function in older adults [15, 16]. However, significant inter-individual variability exists with respect to the influence of increased brain Aβ concentrations on cognitive and motoric outcomes. Hence, identifying common modifiable protective factors that could attenuate age- and disease-related declines in cognitive and physical function is paramount. Specifically, identifying factors that explain inter-individual variability in associations between Aβ and clinical outcomes among non-demented older adults could inform interventions designed to delay declines in both cognitive and physical domains of function.

Cognitive reserve (CR) is considered a buffer that confers individual differences in adapting to and executing tasks in the context of adverse effects of aging, disease, or injury to brain structure and function [17]. Extensive research [18, 19] including a recent meta-analysis [20] demonstrate robust protective effects for CR against cognitive decline and dementia. In contrast, literature concerning the role of CR in mitigating age-related effects on physical outcomes is scarce. Recent studies, however, demonstrated that, among community-residing older adults, higher CR is associated with more efficient brain activation patterns implicated in single and dual-task walking [21], and lower odds of developing incident mobility impairment [22, 23]. Combined, these findings suggest a more generalized role for CR in mitigating risk of developing adverse health outcomes wherein individuals with higher CR levels are less susceptible to both cognitive and motoric declines.

The assessment of CR, however, is controversial as various proxy measures and methods have been utilized to operationalize this hypothetical construct [19]. One approach is to identify measures of brain network connectivity that correlate with CR or modify the associations between neuropathological measures and clinical outcomes. Using functional magnetic resonance imaging (fMRI) to assess resting-state blood oxygenation level dependent (BOLD) [24] to characterize brain network properties has the advantage that the measures are task-invariant and have been shown to be replicable across studies, populations, and neuroimaging modalities [25]. Indeed, increased resting-state connectivity, notably in frontal regions, is associated with CR, and modifies the associations between cortical thickness and cognitive function [26]. Converging evidence further suggests that increased functional connectivity at rest in executive frontal networks may be considered a functional neural marker of CR [27-29]. In addition, increased functional organization of brain networks is associated with higher socio-demographic and performance measures of CR [30], and attenuates the negative effect of neuropathological markers of Alzheimer’s disease on cognitive decline [31]. The strength of functional network connectivity in an episodic memory network mediated the relationship between brain tau levels measured using positron emission tomography (PET) and episodic memory above what as accounted for by cortical thickness in cognitively unimpaired adults [32]. This finding does not imply CR *per se* but does indicate that the network connectivity is the neural mechanism linking pathology measures to cognition. In sum, there is conceptual and empirical support to using resting-state brain networks as a neural surrogate for CR. However, whether inter-individual differences in resting-state network measures of CR explain variability in associations between Aβ and cognitive and motor outcomes in healthy older adults has not been reported.

The current study addressed this critical gap in the literature. Specifically, we evaluated whether functional brain network measures of CR interacted with brain Aβ (measured using positron emission tomography [PET]) in associations with baseline and longitudinal measures of cognitive and physical function. We used data from the Brain Networks and Mobility (B-NET) study [33], a longitudinal study to determine whether brain network community structure [34] could shed light on cortical involvement in cognitive and physical function and decline among older adults. We employed an un-biased empirical approach to operationalize neural representation of CR using brain network community structure [34] in eight canonical resting-state networks commonly identified across different methods [35-42]. These intrinsic networks included the central executive network (CEN), basal ganglia network (BGN), default mode network (DMN), sensorimotor network (SMN), dorsal attention network (DAN), salience network (SN), frontotemporal network (FTN), and visual network (VN). The analysis assigned groups of network nodes into a community if they are more connected with each other than with other network communities. We then quantified the spatial alignment of each participant’s community structure with the eight intrinsic networks [43] as a measure of network integrity. We hypothesized that the integrity of the network community structure would statistically interact with cross-sectional and longitudinal associations of Aβ with cognitive and physical function. Specifically, we predicted that higher CR (i.e., greater intrinsic network integrity) would attenuate negative associations between Aβ concentrations and cognitive and physical function. The digit symbol substitution test (DSST) [44] and expanded short physical performance battery (eSPPB) [45] were used as measures of cognitive and physical function, respectively, as it has been proposed that they share common neural mechanisms [46].

## 2. Methods

### 2.1 Participants

B-NET is a longitudinal, observational trial of community-dwelling older adults aged 70 and older recruited from Forsyth County, NC that enrolled 192 participants. Participants were asked to complete a baseline visit along with follow-up visits at 6, 18, and 30 months. Brain MRIs that were collected at baseline and cognitive and physical function scores from baseline and 30 months were used in the current analyses. The current study focuses on 81 participants that participated in an ancillary study that included positron emission tomography (PET) imaging. All participants gave written informed consent in this study as approved by the Wake Forest University School of Medicine Institutional Review Board (IRB, protocol #IRB00046460).

The following were exclusion criteria for the parent study: hospitalization or surgery within the past 6 months, uncontrolled or serious chronic disease, uncorrected major hearing or vision problems, single or double amputee, musculoskeletal implants physical functional testing (e.g., joint replacements), dependency on assistance for ambulation, clinical diagnosis of any disease affecting mobility (e.g., Parkinson’s disease), psychotic disorder, alcohol use disorder, history of traumatic brain injury or brain tumor, recent history of seizures, or a score of 20 or lower on the Montreal Cognitive Assessment (MoCA). MOCA scores from 21-25 along with other cognitive test results were reviewed by the study neuropsychologist to determine eligibility on an individual basis. Other exclusion criteria included the inability/unwillingness to complete a brain MRI or PET scan, plans to move from the area within 24 months, or current participation in a behavioral intervention trial.

### 2.2. Cognitive and physical function assessments

The Digit-Symbol Substitution Test (DSST) [44] was used to assess cognitive processing speed. This is a paper and pencil test where digits 1-9 are associated with a symbol. The digit-symbol assignments are displayed at the top of the page. Below that are rows of digits with empty boxes below each digit. The participants are instructed to copy the symbol associated with each digit. They are first given 7 practice items and then instructed to fill in as many of the remaining boxes as they can in two minutes. The total number of correct symbols is the final score that was used in the analyses.

The Expanded Short Physical Performance Battery (eSPPB) [45] was used to assess overall physical function. This test was adapted from the original SPPB [47] to be used in well-functioning populations. The four components of the eSPPB include balance, four-meter walk, narrow four-meter walk, and a chair stand. The balance assessment includes a side-by-side posture, semi-tandem, tandem, and one-leg positions. For the 4-meter walk participants are timed walking at their usual speed for two trials and the fasted is used. For the “narrow walk” participants are given 3 attempts to complete two successful walks not stepping outside of a 20 cm path. For the chair stand, participants were timed while standing up from a seated position 5 times without using their arms. Scores for each test within the eSPPB assessment ranged from 0-1 based on a ratio of the measured value to the best possible performance. Adding across the four components gives a continuous score from 0-4.

### 2.3 Aβ PET

PET scans began approximately one year after the parent study with the average time between the participant’s baseline visit and the subsequent PET scan being 260.64 days. PET scans were not completed at any of the follow-up visits. [11C]PiB was used to assess fibrillar Aβ brain deposition on PET. Following a CT scan for attenuation correction, participants were injected with ∼370 MBq [11C]PiB and scanned on a 64-slice GE Discovery MI DR PET/CT [48]. Centiloid (CL) analysis [49] was conducted in PMOD v4.1 (PMOD LLC Technologies, Switzerland). PET frames were aligned to a 3D T1-weighted MRI, and a static PET image was created by averaging 50-70 min (5-min frames) post injection data. The MRI and aligned average PET scan were input into the PMOD PNEURO Step-wise Maximum Probability Atlas workflow using the CL atlas template. MRIs were normalized to MNI-space template and segmented, with coregistered PET scans normalized to MNI space using MRI parameters. SUVr was calculated using the standard MNI-space CL ROI (whole cerebellum reference), and CL scores were calculated using Klunk et al. [49] equation 1.3b (CL =100(SUVr-1.009)/1.067). This method was validated using the GAAIN data set. All reports of Aβ levels are the whole brain quantitative CL scores. All analyses used CL scores as a continuous measure, but for descriptive purposes a cut-off of >24 was used to identify positive scans [50].

### 2.4. MRI Collection, Processing, and Network Generation

All participants completed brain MRI scans at baseline and at the 30-month follow-up. The current study used the baseline MRI scans. Scans included a T1-weighted 3D volumetric MPRAGE anatomical image (TR=2300ms; TE=2.98ms; number of slices=192; voxel dimensions = 1.0×1.0×1.0mm; FOV=256mm; scan duration=312s), T2 FLAIR images (TR=4800ms; TE=4.41ms; number of slices=160; voxel dimensions=1.0×1.0×1.2mm; FOV=256mm; scan duration=293s), and resting-state blood oxygenation level-dependent (BOLD) images (TR=2000ms; TE=25ms; number of slices=35; voxel dimensions =4.0×4.0×5.0mm; FOV=256mm; scan duration=7m 20s). During the resting-state fMRI scan, a fixation cross was displayed on the monitor. Other image sequences were performed but are not used in the current study.

Images were preprocessed using Statistical Parametric Mapping version 12 (SPM12, http://www.fil.ion.ucl.ac.uk/spm), FMRIB’s “topup” Software Library (FMRIB Software Library v6.0) and Advanced Normalization Tools (ANTs). Structural images were segmented based on gray and white matter and cerebrospinal fluid using SPM12. Gray and white matter segmented images were then summed to generate a mask of brain parenchyma. The summed images were manually cleaned to removed extra-parenchyma tissues using MRIcron software [51]. Images were then masked and spatially normalized according to the Montreal Neurological Institute (MNI) template using ANTs.

Functional image preprocessing included dropping first 10 image volumes, fieldmap distortion correction, slice time correction, realignment, coregistration with native-space anatomical images, and warping to MNI space using transformation information from ANTs. Signals from total white matter, total gray matter, total CSF, and the 6 rigid-body motion parameters were regressed from the functional images and the data were band pass filtered (0.009-0.08Hz). The motion scrubbing procedure developed by Power and colleagues [52] was used to correct head motion artifacts during the scan. Further details of structural and functional image processing including the calculation of structural brain image covariates are in the supplemental Methods.

### 2.5 Brain Network Analyses

Networks were generated by performing voxel-wise cross-correlations on each voxel pair. The resulting matrix is a weighted brain network, where each voxel is a node and correlation coefficients between nodes are edges. An empirically determined threshold was calculated to satisfy the equation S=log(N)/log(K), where N is the number of network nodes (∼20,000), K= average number of connections per node, and S was set to 2.5 based on prior research [53]. Correlation coefficients at or above the threshold were set to 1 indicating the presence of a connection, and those below the threshold were set to 0. The threshold was applied to the matrix to dichotomize the data and create a final N x N binary adjacency matrix, Aij. This procedure ensures networks with comparable density across participants.

Modularity (Q) [34] was used to identify network community partitions for each study participant in each condition using a dynamic Markov process [54]. The partitioning procedure resulted in each individual participant’s brain network being divided into categorical communities. Given the exploratory nature of the study, a data-driven approach was used where each participant’s communities were compared to *a priori* templates for eight subnetworks that cover the entire brain using a spatial similarity index called scaled inclusivity, or SI [55]. The subnetworks included the central executive network (CEN), basal ganglia network (BGN), default mode network (DMN), sensorimotor network (SMN), dorsal attention network (DAN), salience network (SN), frontotemporal network (FTN), and visual network (VN). The resulting SI values were used as a measure of brain network structure in the regression analyses detailed below. Further information on the SI analyses can be found in the supplemental Methods section.

### 2.5 Statistical Analyses

Baseline descriptive statistics (i.e., mean, standard deviation, proportions) were calculated separately for the participants in the PET cohort and the remainder of the BNET participants. T-tests were used to compare means between groups for continuous variables, and chi-square tests were used for proportions. All data plots used in figures were generated with Prism (https://www.graphpad.com). All baseline data are represented with filled circles and 30-month data with filled triangles.

Regression analyses were performed using cognitive (DSST) and physical function (eSPPB) measures as outcome variables to identify associations with brain predictors as independent variables. The regression analyses used distances that were calculated with a 3-dimensional SI brain map of the network community structure for each participant. Distance was computed between every subject pair to create a distance matrix for each independent variable. The Jaccard distance (see supplemental Methods section) was to quantify the distance between the SI community structure maps. Absolute distance was used for all continuous variables, including the outcome variables. A linear statistical model with individual-level effects [56] was used to regress physical and cognitive measure distances against predictor variable distances. Further information on the statistical analyses can be found in the supplemental Methods section.

Separate analyses were performed using baseline and 30-month assessments of DSST and eSPPB. Pearson’s correlations were used to assess associations between whole-brain Aβ levels and cognitive (DSST) and physical function (eSPPB). To determine if network community structure modified the association between Aβ and DSST or eSPPB, distance regression analyses [56], as described above, were used. For the primary analyses, models included the main effects for the network community structure, Aβ, and the network community structure *Aβ interaction to assess if community structure modifies the relationship between Aβ and the outcome measures. Models were run for each of the 8 intrinsic networks. This method allows for the identification of associations between brain network spatial patterns and other numeric or categorical variables. No covariates were included in the primary, unadjusted analyses. An adaptive false discovery rate was applied to correct for multiple comparisons across all primary analyses [57, 58] and the resultant q-values were used to for assessing statistical significance (q < 0.05). Only brain networks that remained statistically significant after correction for multiple comparisons were used in secondary, adjusted analyses with additional covariates. These further analyses report nominal p-values.

In addition to the main analyses, three different sets of supplemental analyses were performed for the models that exhibited a significant main or interaction effect. All 3 of the analyses were adjusted for additional covariates to control for participant age and for participant head motion in the MRI scanner. One set of analyses included the time between the completion of the main baseline study visits and the PET imaging session. Because the PET was an ancillary study, the PET scans did not begin until after the study was ongoing. These analyses were intended to ensure that differences in the delay to get the PET scan was not having a major effect on the outcomes. Another set of analyses was performed to include structural brain measures that are commonly used in studies of cognitive reserve. The intent of this set of analyses was to determine if the brain network* Aβ interactions remained significant while accounting for age-related changes in brain structure. The third set of analyses was a sensitivity analysis performed to address the fact that each of the analyses had a different number of participants due to different missing data. 76 of the total 81 participants were used in these analyses as they all had DSST and eSPPB scores at baseline and at 30 months.

## 3. Results

### 3.1 Participants

The demographic details of the study participants are presented in Table 1. The measures for the PET cohort were compared to the remainder of the study participants that were not in the PET cohort. There were no differences between groups for the variables assessed, indicating that the PET sample is representative of the groups as a whole.

**Table 1.** Baseline Characteristics of Participants By PET Cohort.

|  | *Not in PET Cohort*<br>*(n=111)* | *In PET Cohort*<br>*(n=81)* | *Overall*<br>*(n=192)* | *P-value* |
| --- | --- | --- | --- | --- |
| **Age** | **76.43 (4.72)** | **76.43 (4.75)** | **76.43 (4.72)** | **0.9970** |
| **Race/Ethnicity** |  |  |  | **0.5689** |
| White | 98 (88.3) | 73 (90.1) | 171 (89.1) |  |
| African American/Black | 12 (10.8) | 6 ( 7.4) | 18 ( 9.4) |  |
| American Indian or Alaskan Native | 1 ( 0.9) | 1 ( 1.2) | 2 ( 1.0) |  |
| Asian | 0 ( 0.0) | 1 ( 1.2) | 1 ( 0.5) |  |
| **Sex** |  |  |  | **0.1789** |
| Men | 44 (39.6) | 40 (49.4) | 84 (43.8) |  |
| Women | 67 (60.4) | 41 (50.6) | 108 (56.3) |  |
| **Years of Education** | **15.67 (2.20)** | **15.70 (2.77)** | **15.68 (2.45)** | **0.9178** |
| **PET measures** |  |  |  |  |
| Days from last Baseline Visit to PET Visit |  | 260.64 (112.42) |  |  |
| Centiloid Score |  | 25.39 (40.29) |  |  |
| # positive scans (CL > 24) |  | 29 |  |  |
| **MRI measures** |  |  |  |  |
| Total gray matter volume in CC | 596.12 (87.05) | 606.16 (70.97) | 600.36 (80.61) | 0.3958 |
| Bilateral hippocampus volume in CC | 7.15 (0.91) | 7.31 (0.99) | 7.22 (0.95) | 0.2658 |
| Bilateral thalamus volume in CC | 4.60 (0.81) | 4.70 (0.92) | 4.64 (0.85) | 0.4442 |
| Bilateral prefrontal cortex volume in CC | 31.51 (4.72) | 31.88 (5.20) | 31.67 (4.92) | 0.6054 |
| **MoCA Adjusted Score** | **25.68 (2.19)** | **25.58 (2.22)** | **25.64 (2.20)** | **0.7672** |
| **DSST** | **55.57 (12.62)** | **54.65 (11.65)** | **55.18 (12.20)** | **0.6097** |
| **eSPPB** | **2.00 (0.53)** | **2.01 (0.52) (n=79)** | **2.00 (0.52)** | **0.8712** |

### 3.2 Cognitive and Physical Function Scores

Summary statistics of performance on the DSST and eSPPB are shown in Table 1. At baseline, the population had an average DSST score of 54.7 and an average eSPPB score of 2.01. The population means did not change meaningfully at 30 months, with an average DSST score of 55.4 and an average eSPPB score of 2.01. The distributions for both measures were normal at baseline (DSST: KS = 0.08, p>0.1, eSPPB: KS = 0.07, p>0.1) and at 30 months (30m DSST: KS = 0.07, p>0.1, 30m eSPPB: KS = 0.09, p>0.1).

### 3.3 Aβ Associations with Cognitive and Physical Function

Should high brain amyloid be associated with assessments of cognitive and/or physical function, it would be expected that those individuals that are Aβ positive would have the poorest scores. In this population, such a pattern was not seen at baseline or at 30 months (Figure 1). It is clear from the figure that Aβ positive individuals are represented throughout the distribution of scores for both DSST and eSPPB at both time points. In fact, the Aβ positive group had an average DSST and eSPPB within 1 standard deviation of the mean for the entire population at baseline (DSST:53.9, eSPPB: 1.88) and 30 months (DSST:51.9, eSPPB: 1.88).

**Figure 1.**
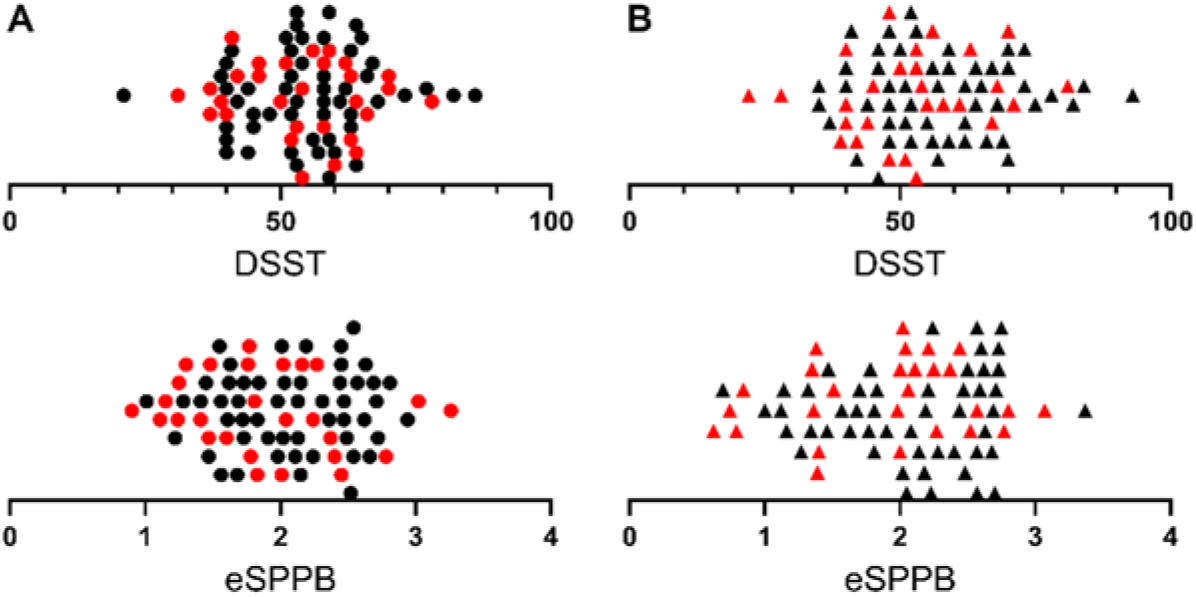
Distribution of participant DSST and eSPPB scores at baselin (A)and 30 months (B). Participants with positive Aβ scans are red. Note the Aβ positive individuals are concentrated at the lower ends of the distribution but are represented throughout the score ranges at baseline and 30 months. Individual participant data is represented by circles at baseline and triangles at 30 months. Data points were algorithmically spread vertically for visualization purposes.

Correlations between whole-brain Aβ levels and eSPPB and DSST revealed no significant association at baseline for either measure (Figure 2A). There was a slight negative slope for eSPPB (i.e., higher Aβ may be related to lower eSPPB scores) that didn’t reach significance; we note that most of the individuals with Aβ levels near 100 CL had eSPPB scores close to the group mean. An individual with CL = 94 actually had the highest eSPPB score of the population. At the 30-month follow-up (Figure 2B), the relationship between DSST and Aβ levels had a slight negative slope, but it was not significant. The relationship between eSPPB and Aβ achieved significance at 30 months but was still relatively weak with a slope of -0.004 and this relationship only accounted for 7% of the variance in eSPPB scores.

**Figure 2.**
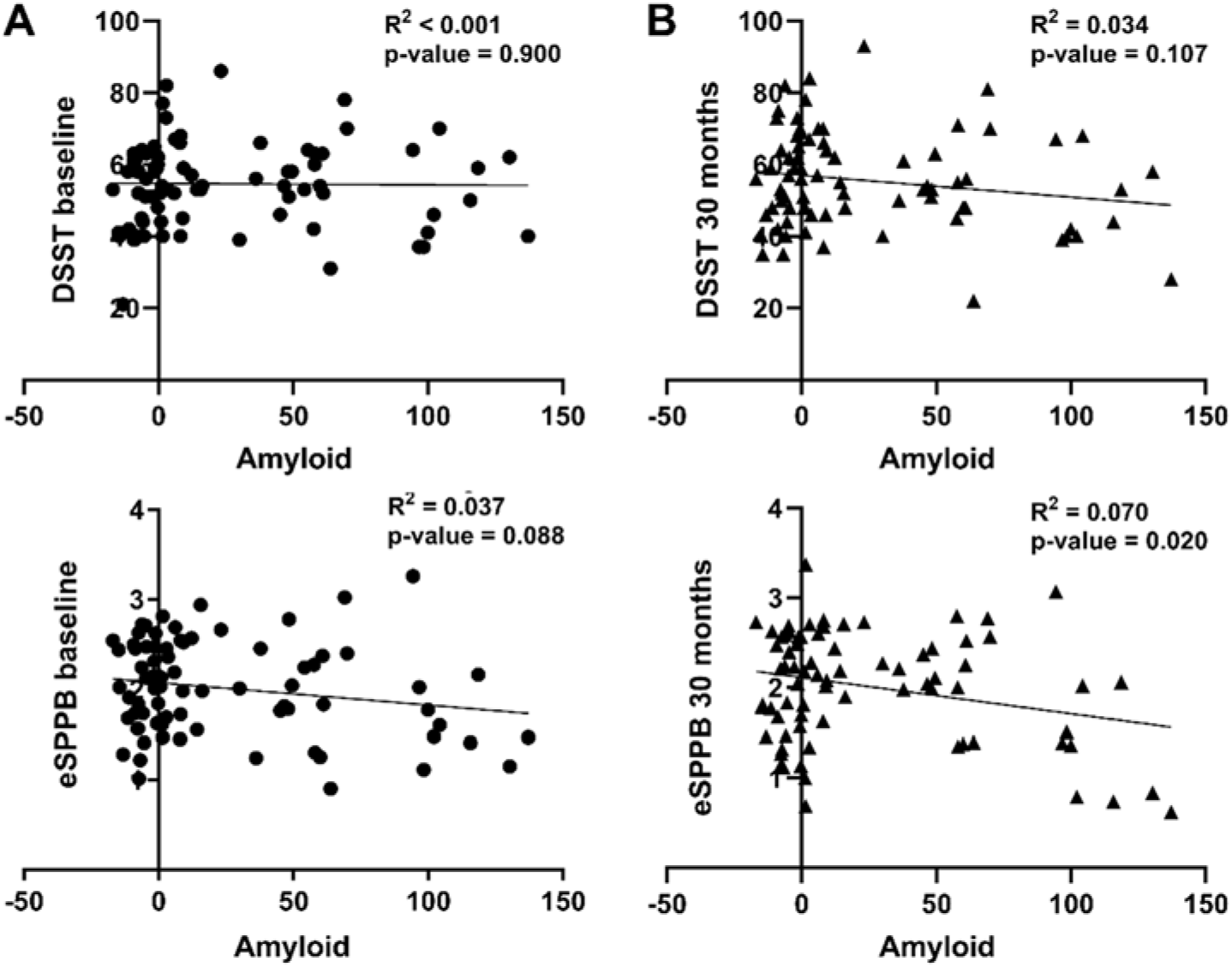
Associations of Aβ with DSST and eSPPB scores at baseline (A) and 30 months (B). Due to missing data, there are different numbers of participants in each analysis. For DSST there were 81 at baseline and 78 at 30 months. For eSPPB there were 79 at baseline and 77 at 30 months. Individual participant data is represented by circles at baseline and triangles at 30 months.

### 3.4 Brain network community structure and Aβ modification analyses

Effect modification of community structure of the 8 intrinsic brain networks on the relationships between Aβ levels outcome variables (DSST and eSPPB) were assessed using statistical interactions. The CEN and BGN exhibited significant interactions with Aβ for both DSST and eSPPB with correction for multiple comparisons (Table 2). All estimates for CEN, BGN, and amyloid that are listed in Table 2 are those observed when the interaction was present in the model. The results demonstrate that interactions consistent with cognitive reserve were present at baseline for eSPPB but were not present until the 30-month follow-up for the DSST. For the eSPPB, the CEN and BGN interactions were both significant at baseline with similar effect sizes. The CEN remained significant at 30 months, but the BGN did not achieve significance. For the DSST, there were no significant interactions at baseline. However, the main effects for both the CEN and BGN were significant at baseline. At 30 months the Aβ*network interactions were both significant with comparable effect sizes that were ∼3 times greater than effects observed at baseline. For completeness, results for the networks that did not exhibit significant main or interaction effects are in the supplement materials (Table S1).

**Table 2.** Regression results for the CEN and BGN.

| Outcome Variable | Independent Variables | Estimate | SE | T score | p-Value | FDR q-Value |
| --- | --- | --- | --- | --- | --- | --- |
| Baseline eSPPB | amyloid | -0.0066 | 0.0020 | -3.2953 | 0.0010 | 0.0127 |
|  | CEN | 0.8278 | 0.2552 | 3.2442 | 0.0012 | 0.0127 |
|  | **amyloid\*CEN** | **0.0099** | **0.0028** | **3.4911** | **0.0005** | **0.0127** |
|  | amyloid | -0.0072 | 0.0023 | -3.0875 | 0.0020 | 0.0187 |
|  | BGN | 0.1286 | 0.2650 | 0.4851 | 0.6276 | 0.8248 |
|  | **amyloid\*BGN** | **0.0098** | **0.0030** | **3.2489** | **0.0012** | **0.0127** |
| Baseline DSST | amyloid | -0.0516 | 0.0403 | -1.2806 | 0.2004 | 0.4487 |
|  | **CEN** | **24.7596** | **5.1641** | **4.7946** | **0.0000** | **0.0002** |
|  | amyloid*CEN | 0.0697 | 0.0571 | 1.2204 | 0.2224 | 0.4604 |
|  | amyloid | -0.0559 | 0.0465 | -1.2026 | 0.2292 | 0.4604 |
|  | **BGN** | **16.0647** | **5.2288** | **3.0724** | **0.0021** | **0.0187** |
|  | amyloid*BGN | 0.0695 | 0.0606 | 1.1475 | 0.2513 | 0.4762 |
| 30-month eSPPB | amyloid | -0.0082 | 0.0024 | -3.4125 | 0.0007 | 0.0127 |
|  | CEN | 0.4014 | 0.3141 | 1.2777 | 0.2015 | 0.4487 |
|  | **amyloid\*CEN** | **0.0127** | **0.0034** | **3.7287** | **0.0002** | **0.0094** |
|  | amyloid | -0.0057 | 0.0028 | -2.0233 | 0.0431 | 0.1532 |
|  | BGN | 0.0555 | 0.3255 | 0.1704 | 0.8647 | 0.9697 |
|  | amyloid*BGN | 0.0084 | 0.0037 | 2.2801 | 0.0227 | 0.1090 |
| 30-month DSST | amyloid | -0.1596 | 0.0486 | -3.2854 | 0.0010 | 0.0127 |
|  | CEN | 18.9454 | 6.3188 | 2.9983 | 0.0027 | 0.0219 |
|  | **amyloid\*CEN** | **0.2311** | **0.0687** | **3.3634** | **0.0008** | **0.0127** |
|  | amyloid | -0.1561 | 0.0567 | -2.7510 | 0.0060 | 0.0410 |
|  | BGN | 16.2112 | 6.5487 | 2.4755 | 0.0134 | 0.0715 |
|  | **amyloid\*BGN** | **0.2072** | **0.0737** | **2.8119** | **0.0050** | **0.0366** |

Significant interactions or main effects in the absence of an interaction are bolded. SE- standard error
CEN - Central Executive Network, BGN - Basal Ganglia Network, FDR - false discovery rate
DSST - digit symbol substitution test, eSPPB - expanded short physical performance battery

In all analyses with significant main or interaction effects, the estimates for the main or interaction effects were positive. This indicates that as the difference between subject brain network patterns increased, the difference in the behavioral outcome increases. However, the direction of the relationships for brain networks and Aβ with the behavioral outcomes cannot be determined directly from the estimates. To better interpret the observed significant interactions, plots were developed to capture the four quadrants of the interactions (high network integrity/low Aβ, high network integrity/high Aβ, low network integrity/low Aβ, low network integrity/high Aβ). The division for Aβ was not simply clinically positive or negative scans as the groupings included high and low network integrity as well. See supplemental methods section for a description of how the individuals in each group were identified. Note that these figures are for visualization purposes as the statistical outcomes were based on a continuous regression including all available data for each outcome variable.

The significant interactions that the CEN and BGN had with Aβ at baseline for eSPPB and at 30 months for DSST are explored graphically in interaction plots (Figure 3). Baseline interactions for the eSPPB showed very similar relationships for the two networks. The mean eSPPB score was very similar for the low and high Aβ groups when the networks had high integrity. When networks had low integrity, the mean eSPPB score was lower with the high Aβ being the lowest. Thus, high levels of Aβ were only associated with lower eSPPB scores if the networks had low integrity. The CEN* Aβ interaction for DSST at 30 months exhibited a similar pattern. Mean DSST score was highest when the networks had high integrity, with the low Aβ groups having the highest DSST. Low network integrity was associated with a lower mean DSST score with the high Aβ group having the lowest score. The BGN * Aβ interaction for DSST at 30 months was somewhat different. As observed with the

**Figure 3.**
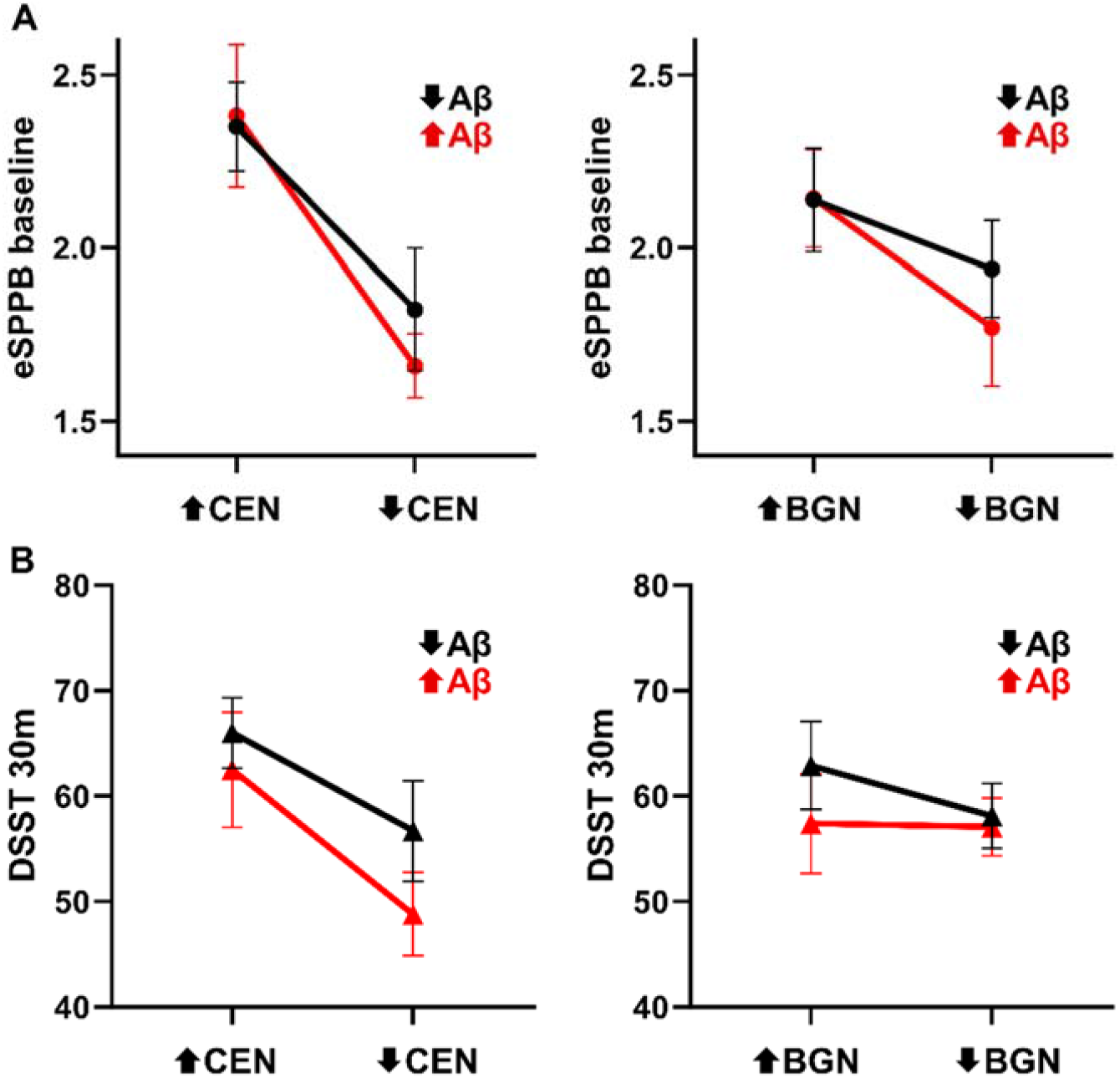
Plots depicting the direction of the significant interactions between brain networks and Aβ for eSPPB at baseline (A) and the DSST at 30 months (B). The data show the mean and the standard error of the mean (SEM) for the four categorical groups. The figure should be interpreted qualitatively to aid in understanding the direction of the relationships between network integrity/Aβ and behavioral test scores. All quantitative assessments should be based on the regression results that used continuous measures.

CEN, the high network integrity/low Aβ group had the highest mean DSST score and the mean DSST score was lower in the low network integrity/low Aβ. Those with high Aβ had a lower mean DSST score than those with low Aβ and there was no notable effect of network integrity. The mean Aβ levels for the high and low groupings are shown in Figure S1. The group average images of the CEN and BGN for the high and low network integrity groups are shown in Figure S2.

### 3.5 Adjusted Analyses

For analyses that included the time between the completion of the baseline visits and the PET scan, age, and head motion during the scan, age was significant in all models. The PET scan date was significant in all models except the 30-month DSST models. Head motion was only significant for the baseline DSST. Despite the significance of the covariates, there were no meaningful changes in effect sizes or significance of the network/Aβ outcomes for these analyses (Table S2). Analyses that included structural brain measures revealed that the effect sizes for the brain network* Aβ interactions were reduced but all remained significant except the BGN* Aβ interaction for the DSST at 30 months (Table S3). The effect size for this interaction was reduced by about ½. It is notable that the main effect of the BGN was significant in this adjusted analysis (this indicates that when Aβ is equal to zero CL the BGN was significantly associated with DSST score). In addition, there were multiple significant associations between brain anatomy variables and the behavioral measures. As these outcomes were not the primary of objective of these analyses, the significant findings are not detailed here but can be found in Table S3. A sensitivity analyses that used the same 76 participants across all models to account for missing data also included age, PET date, and head motion as covariates. There were only minimal changes in effect sizes and no change in significance in the analyses (Table S4).

## 4. Discussion

The current study utilized resting state functional brain networks to operationalize CR and determined their utility in modifying baseline and longitudinal associations of Aβ with cognitive and physical outcomes in older adults free of dementia at baseline. Consistent with the study hypotheses, we found that the integrity of the central executive and basal ganglia networks modified associations of Aβ with cognitive and physical performance. These modification effects, however, varied as a function of study outcome and time. For the physical function measure, the modification was present at baseline, but the modification was not present until the 30-month follow-up for the cognitive measure. In all cases, the CEN exhibited stronger modification than the BGN. These findings suggest that CR, as defined herein, may have protective effects against Aβ that extend to both cognitive and motor outcomes. A nuanced alternative interpretation is that resting-state network integrity represents a broader reserve buffer than CR and may thus confer protective effects generalizable to multiple domains of function in aging and disease. These findings are discussed below.

Prior work has shown that functional connectivity in fronto-striatal circuits [59] including the central executive [60-62] and basal ganglia [63, 64] networks is associated with cognition in aging. We found that, at baseline, higher integrity of both networks was associated with better DSST performance, a measure of speed of processing and working memory. There were no associations between Aβ and

DSST at baseline. For cognitive performance at the 30-month follow-up, there were significant two-way interactions wherein higher integrity in both networks was protective against higher Aβ concentrations. These findings provide first evidence that inter-individual differences in CR, operationalized via measures of intrinsic functional brain network integrity, modifies associations of Aβ with future cognition in dementia-free older adults. Discrepancies between cross-sectional and longitudinal associations are common [65]; and in the current study non-significant modification effects (i.e., interactions) between network connectivity and Aβ at baseline may be attributed to reduced variance on DSST performance given the healthy nature of this sample.

Literature concerning cognitive [8-10] and brain [66-68] control of physical function in aging is robust. However, knowledge concerning the role of resting state networks in physical function is relatively scarce in older adults [33]. Notably, a recent study revealed that connectivity in the frontoparietal network is implicated in motor skill learning [69]. Our study extends previous findings providing first evidence that integrity in both the central executive and basal ganglia networks modified associations between Aβ and eSPPB performance at baseline. The association between Aβ and eSPPB performance at 30-month follow-up was modified only by the central executive network. This suggests that higher integrity in a network implicated in higher order cognitive processing served as a buffer against poor current and future physical performance. Recent studies indicate that higher CR is related to more efficient brain control of walking, notably under attention-demanding conditions [21], and also attenuates the risk of developing incident mobility impairment [22, 23]. Hence, building on the extant limited literature, the current study introduces novel evidence for the role of CR, defined using connectivity in the aforementioned networks, as a buffer against the influence of Aβ on physical function.

It may be argued that integrity of the central executive and basal ganglia networks is a proxy for a buffer that is broader than CR. That is, a construct, previously termed individual reserve [70], that captures collective cognitive and physical reserve capacities within a person may better account for the ability of the brain to adapt to cognitive and physical consequences of aging. This view of reserve provides an additional lens through which cognitive-motor interactions may be understood in the context of aging and age-related neurodegeneration. However, the fact that we did not see any significant interactions between Aβ and other networks, such as the DMN, suggests that the findings are network specific.

This study is not without limitations. The study was a secondary analysis using a subset of older adult participants from the larger, parent B-NET study. The participants in the parent BNET study were specifically recruited to have normal cognition at baseline as the primary objective was to examine brain-motor associations. It is possible that the findings observed here are specific to those individuals that exhibit normal cognition. This would remain an important finding as it could be evidence for neurobiological mechanisms that prevent decline. Although to our knowledge this is the first study to combine brain networks and amyloid in examining associations with cognitive and physical function, the study sample is relatively small. Furthermore, the sample is fairly homogenous, and future studies should not only expand the sample size but should also strive to include a more diverse population.

The distance regression methodology employed here has been validated [56] and used to evaluate brain network-behavior associations [33]. However, associations between spatial brain patterns and behavioral measures are inherently complicated, and the regression estimates can be difficult to interpret. For this reason, a single “global” average CL score was used rather than using the Aβ images. Future work could expand on these findings by including the spatial Aβ patterns or CL scores for each brain subnetwork. Finally, the field of brain network analyses is rapidly evolving and there is no standardized methodology that is universally accepted. We have chosen to use image processing and network analysis methods that we have found robust in our prior work on older adults [71-75].

Replication of our results using other study data and processing methods will help demonstrate the robustness and generalizability of the findings.

### Clinical Implications and future direction

These findings are notable given that Aβ-targeting pharmacological agents are quite effective at lowering brain amyloid levels but have small [76-79] or no [80] effect on cognitive function and have significant, potentially life-threatening side effects [81]. It has also been shown that older adults with elevated brain Aβ deposition in their brains can have normal cognition [82, 83], as was found in more than 35% of the participants in this study. There are likely heterogenous responses to the Aβ reducing drugs and it may be that a major underlying factor is brain network integrity. Given that the current study was limited to persons with normal cognition, it is not known how Aβ and brain networks may interact in those with cognitive impairment. Future studies could investigate the relationships found here in individuals with cognitive impairment and could consider analyses examining apolipoprotein E (APOE) 4 carriers and noncarriers separately or include the genotype as an interacting factor. If future research confirms that brain network integrity and Aβ interact not only in normal cognition but also in cognitive decline, then it may be pertinent to assess network integrity in individuals prior to the initiation of amyloid lowering treatments.

## Supporting information

Supplemental methods and result

## Acknowledgements

This work was supported by the National Institute on Aging (AG052419, AG052419-05S1, and P30 AG021332); the National Institute of Biomedical Imaging and Bioengineering (EB032903); and the National Center for Advancing Translational Sciences (UL1TR001420). We would like to thank the entire BNET study team that contributed to the participant recruitment, data collection, and data curation/analyses. We would also like to thank the BNET participants that contributed substantial amounts of time and effort toward participation.

