## Supplemental methods and result for "Resting-state connectivity modifies the effects of amyloid on cognitive and physical function: evidence for network-based cognitive reserve"

### 1. Supplemental Methods

#### 1.1. Image analyses

The T1 weighted images were segmented into grey matter (GM) and white matter (WM) and cerebrospinal fluid (CSF) using SPM-12. The resulting GM and WM segments were used to calculated GM volume and WM volume respectively. Intracranial Volume (ICV) was calculated by combining all three (GM, WM, and CSF) segments. White matter lesions were calculated using the Lesion Segmentation Toolbox LPA with default settings. Not threshold was applied resulting in each voxel containing a value that represents the percentage of the voxel that is a lesion. The output image was used to calculate white matter lesion volume (WMLV). Volumetric measures based on segmented images (GMV, WMV, WMLV, ICV) are in cubic centimeters and were calculated by summing all voxels values from the appropriate image. For example, for GMV the gray matter volume image was used with each value indicating the volume of gray matter in that voxel. Images were then multiplied by the cubic volume of the voxel and dividing by 10^3 resulting in all measures being in cubic centimeters (CC). To calculate the volume of structure brain regions (hippocampus, thalamus, prefrontal cortex), the appropriate brain atlases were warped to each subject's native space to calculate accurate volumes. The volume of each structure was determined in cubic centimeters. The Automated anatomical labelling atlas 2 (AAL2) [1] was used to calculate volume of hippocampus (regions 41 and 42) and thalamus (regions 81 and 82) volumes. The prefrontal cortex volume was calculated with regions 5, 6 and 7 from the Sallet Dorsal Frontal Parcellation [2].

To demonstrate the relationships that underlie the significant distance regression moderation effects, we examined the data to evaluate the dependent outcomes (eSPPB/DSST scores) that were related to high/low amyloid and high/low network integrity. Four groups of individuals were identified for each of the network analyses (CEN and BGN): high network integrity/low Aβ, high network integrity/high Aβ, low network integrity/low Aβ, low network integrity/high Aβ. To rank the community structure, the Euclidean distance between each participant's SI_ROI image for CEN/BGN and the CEN/BGN network template was computed. Individuals with better community structure would be expected to have a community that more closely matched the a priori template. The top and bottom 1/4 participants based on network integrity rank were selected. Within the top and bottom network ranking groups, the data were median split based on amyloid centiloid score. This yielded the four grouping for each network. For the CEN there were 10 participants in each of the four groups. For the BGN the groups were not perfectly balanced with 9 ⬆BGN/⬇Aβ, 10 ⬆BGN/⬆Aβ, 8 ⬇BGN/⬇Aβ, and 12 ⬇BGN/⬆Aβ participants. It is important to stress that these groupings were not used for statistical analysis but were used to visually identify the direction of the relationship found in the distance regressions.

Table S1. Model results for networks without significant findings

| Outcome Variable | Independent Variables | Estimate | SE | T score | FDR p-Value | Outcome Variable | Independent Variables | Estimate | SE | T score | FDR p-Value |
|---|---|---|---|---|---|---|---|---|---|---|---|
| DAN | | | | | | | | | | | |
| Baseline DSST | amyloid | -0.0858 | 0.0597 | -1.4375 | 0.3918 | 30-month DSST | amyloid | -0.0850 | 0.0711 | -1.1941 | 0.4604 |
| | DAN | -2.3657 | 5.0782 | -0.4659 | 0.8248 | | DAN | 0.0238 | 6.1144 | 0.0039 | 0.9980 |
| | amyloid*DAN | 0.1062 | 0.0763 | 1.3912 | 0.4143 | | amyloid*DAN | 0.1120 | 0.0909 | 1.2316 | 0.4604 |
| Baseline eSPPB | amyloid | 0.0005 | 0.0030 | 0.1575 | 0.9697 | 30-month eSPPB | amyloid | -0.0007 | 0.0035 | -0.2068 | 0.9697 |
| | DAN | -0.1628 | 0.2520 | -0.6461 | 0.7290 | | DAN | -0.3087 | 0.3084 | -1.0009 | 0.5339 |
| | amyloid*DAN | -0.0002 | 0.0038 | -0.0457 | 0.9980 | | amyloid*DAN | 0.0018 | 0.0045 | 0.4025 | 0.8455 |
| DMN | | | | | | | | | | | |
| Baseline DSST | amyloid | -0.0864 | 0.0540 | -1.6003 | 0.3300 | 30-month DSST | amyloid | -0.1357 | 0.0654 | -2.0751 | 0.1407 |
| | DMN | 6.8118 | 5.2159 | 1.3060 | 0.4487 | | DMN | 3.5836 | 6.3282 | 0.5663 | 0.7721 |
| | amyloid*DMN | 0.1121 | 0.0722 | 1.5524 | 0.3417 | | amyloid*DMN | 0.1852 | 0.0874 | 2.1196 | 0.1363 |
| Baseline eSPPB | amyloid | -0.0042 | 0.0027 | -1.5628 | 0.3417 | 30-month eSPPB | amyloid | -0.0035 | 0.0032 | -1.0859 | 0.5132 |
| | DMN | -0.1649 | 0.2587 | -0.6376 | 0.7290 | | DMN | -0.1911 | 0.3135 | -0.6096 | 0.7433 |
| | amyloid*DMN | 0.0061 | 0.0036 | 1.6935 | 0.2896 | | amyloid*DMN | 0.0056 | 0.0043 | 1.3039 | 0.4487 |
| FTN | | | | | | | | | | | |
| Baseline DSST | amyloid | -0.1243 | 0.0578 | -2.1512 | 0.1363 | 30-month DSST | amyloid | -0.1778 | 0.0687 | -2.5898 | 0.0545 |
| | FTN | 10.8537 | 6.5752 | 1.6507 | 0.3063 | | FTN | -7.1411 | 7.8908 | -0.9050 | 0.5608 |
| | amyloid*FTN | 0.1561 | 0.0742 | 2.1031 | 0.1363 | | amyloid*FTN | 0.2320 | 0.0881 | 2.6330 | 0.0529 |
| Baseline eSPPB | amyloid | 0.0004 | 0.0029 | 0.1315 | 0.9697 | 30-month eSPPB | amyloid | -0.0006 | 0.0034 | -0.1632 | 0.9697 |
| | FTN | -0.0048 | 0.3269 | -0.0147 | 0.9980 | | FTN | -0.1778 | 0.3948 | -0.4503 | 0.8248 |
| | amyloid*FTN | -0.0001 | 0.0037 | -0.0161 | 0.9980 | | amyloid*FTN | 0.0016 | 0.0044 | 0.3672 | 0.8677 |

DAN - dorsal attention network, DMN - default mode network, FTN - frontotemporal network, SMN - sensorimotor network, SN - salience network, VN - visual network

DSST - digit symbol substitution test, eSPPB - extended short physical performance battery, SE - standard error, FDR - false discovery rate

Table S1 cont. Model results for networks without significant findings

| Outcome Variable | Independent Variables | Estimate | SE | T score | FDR p-Value | Outcome Variable | Independent Variables | Estimate | SE | T score | FDR p-Value |
|---|---|---|---|---|---|---|---|---|---|---|---|
| SMN | | | | | | | | | | | |
| Baseline DSST | amyloid | -0.0132 | 0.0540 | -0.2440 | 0.9684 | 30-month DSST | amyloid | -0.0600 | 0.0652 | -0.9204 | 0.5608 |
| | SMN | 1.0788 | 5.5019 | 0.1961 | 0.9697 | | SMN | -6.8361 | 6.6784 | -1.0236 | 0.5339 |
| | amyloid*SMN | 0.0133 | 0.0704 | 0.1886 | 0.9697 | | amyloid*SMN | 0.0816 | 0.0849 | 0.9616 | 0.5472 |
| Baseline eSPPB | amyloid | 0.0000 | 0.0027 | -0.0026 | 0.9980 | 30-month eSPPB | amyloid | 0.0028 | 0.0033 | 0.8717 | 0.5745 |
| | SMN | -0.1219 | 0.2713 | -0.4492 | 0.8248 | | SMN | -0.1419 | 0.3339 | -0.4248 | 0.8366 |
| | amyloid*SMN | 0.0004 | 0.0035 | 0.1275 | 0.9697 | | amyloid*SMN | -0.0028 | 0.0042 | -0.6618 | 0.7279 |
| SN | | | | | | | | | | | |
| Baseline DSST | amyloid | -0.0418 | 0.0435 | -0.9594 | 0.5472 | 30-month DSST | amyloid | -0.1111 | 0.0524 | -2.1210 | 0.1363 |
| | SN | 8.3739 | 5.4680 | 1.5314 | 0.3456 | | SN | -0.0560 | 6.5687 | -0.0085 | 0.9980 |
| | amyloid*SN | 0.0538 | 0.0598 | 0.9002 | 0.5608 | | amyloid*SN | 0.1566 | 0.0718 | 2.1820 | 0.1335 |
| Baseline eSPPB | amyloid | -0.0040 | 0.0022 | -1.8351 | 0.2205 | 30-month eSPPB | amyloid | -0.0062 | 0.0026 | -2.3429 | 0.0970 |
| | SN | -0.3634 | 0.2703 | -1.3446 | 0.4406 | | SN | -0.2806 | 0.3277 | -0.8563 | 0.5790 |
| | amyloid*SN | 0.0059 | 0.0030 | 1.9987 | 0.1567 | | amyloid*SN | 0.0094 | 0.0036 | 2.6213 | 0.0529 |
| VN | | | | | | | | | | | |
| Baseline DSST | amyloid | -0.0555 | 0.0526 | -1.0540 | 0.5289 | 30-month DSST | amyloid | -0.0718 | 0.0627 | -1.1440 | 0.4762 |
| | VN | -4.0989 | 4.3095 | -0.9511 | 0.5472 | | VN | -2.4705 | 5.1611 | -0.4787 | 0.8248 |
| | amyloid*VN | 0.0641 | 0.0641 | 1.0000 | 0.5339 | | amyloid*VN | 0.0907 | 0.0763 | 1.1883 | 0.4604 |
| Baseline eSPPB | amyloid | 0.0000 | 0.0026 | -0.0101 | 0.9980 | 30-month eSPPB | amyloid | -0.0033 | 0.0031 | -1.0365 | 0.5333 |
| | VN | -0.1526 | 0.2148 | -0.7106 | 0.6938 | | VN | -0.3848 | 0.2592 | -1.4846 | 0.3680 |
| | amyloid*VN | 0.0004 | 0.0032 | 0.1363 | 0.9697 | | amyloid*VN | 0.0048 | 0.0038 | 1.2599 | 0.4538 |

DAN - dorsal attention network, DMN - default mode network, FTN - frontotemporal network, SMN - sensorimotor network, SN - salience network, VN - visual network
DSST - digit symbol substitution test, eSPPB - extended short physical performance battery, SE - standard error, FDR - false discovery rate

Table S2. Adjusted models including age, PET date, and head motion

| Outcome Variable | Independent Variables | Estimate | SE | T score | p-Value | Outcome Variable | Independent Variables | Estimate | SE | T score | p-Value |
|---|---|---|---|---|---|---|---|---|---|---|---|
| BGN | | | | | | | | | | | |
| Baseline DSST | amyloid | -0.0420 | 0.0465 | -0.9030 | 0.3666 | 30-month DSST | amyloid | -0.1298 | 0.0565 | -2.2973 | 0.0217 |
| | **age** | **0.1550** | **0.0392** | **3.9491** | **0.0001** | | **age** | **0.3023** | **0.0462** | **6.5471** | **<0.0001** |
| | **PET date** | **0.0070** | **0.0022** | **3.1470** | **0.0017** | | PET date | 0.0050 | 0.0029 | 1.7064 | 0.0880 |
| | head motion | 0.0105 | 0.0223 | 0.4709 | 0.6377 | | head motion | -0.0096 | 0.0263 | -0.3644 | 0.7156 |
| | **BGN** | **15.5554** | **5.2200** | **2.9800** | **0.0029** | | BGN | 15.0364 | 6.5166 | 2.3074 | 0.0211 |
| | amyloid*BGN | 0.0504 | 0.0605 | 0.8323 | 0.4053 | | **amyloid*BGN** | **0.1706** | **0.0734** | **2.3245** | **0.0202** |
| Baseline eSPPB | amyloid | -0.0052 | 0.0023 | -2.2641 | 0.0236 | 30-month eSPPB | amyloid | -0.0039 | 0.0028 | -1.3877 | 0.1653 |
| | **age** | **0.0207** | **0.0019** | **10.8725** | **<0.0001** | | **age** | **0.0196** | **0.0023** | **8.5293** | **<0.0001** |
| | **PET date** | **0.0005** | **0.0001** | **4.3942** | **<0.0001** | | **PET date** | **0.0003** | **0.0001** | **2.0323** | **0.0422** |
| | **head motion** | **0.0027** | **0.0011** | **2.5502** | **0.0108** | | head motion | 0.0016 | 0.0013 | 1.2219 | 0.2218 |
| | BGN | 0.0031 | 0.2597 | 0.0120 | 0.9905 | | BGN | -0.0632 | 0.3221 | -0.1962 | 0.8445 |
| | **amyloid*BGN** | **0.0070** | **0.0030** | **2.3731** | **0.0177** | | amyloid*BGN | 0.0058 | 0.0036 | 1.6054 | 0.1085 |
| CEN | | | | | | | | | | | |
| Baseline DSST | amyloid | -0.0478 | 0.0403 | -1.1870 | 0.2353 | 30-month DSST | amyloid | -0.1444 | 0.0484 | -2.9862 | 0.0028 |
| | **age** | **0.1533** | **0.0390** | **3.9267** | **0.0001** | | **age** | **0.3013** | **0.0460** | **6.5555** | **<0.0001** |
| | **PET date** | **0.0064** | **0.0022** | **2.8885** | **0.0039** | | PET date | 0.0044 | 0.0029 | 1.5147 | 0.1300 |
| | head motion | -0.0022 | 0.0224 | -0.0965 | 0.9232 | | head motion | -0.0236 | 0.0265 | -0.8896 | 0.3737 |
| | **CEN** | **23.6575** | **5.1904** | **4.5579** | **<0.0001** | | CEN | 18.2942 | 6.3466 | 2.8825 | 0.0040 |
| | amyloid*CEN | 0.0634 | 0.0572 | 1.1091 | 0.2675 | | **amyloid*CEN** | **0.2070** | **0.0684** | **3.0246** | **0.0025** |
| Baseline eSPPB | amyloid | -0.0056 | 0.0020 | -2.8728 | 0.0041 | 30-month eSPPB | amyloid | -0.0072 | 0.0024 | -3.0195 | 0.0026 |
| | **age** | **0.0204** | **0.0019** | **10.7800** | **<0.0001** | | **age** | **0.0191** | **0.0023** | **8.3501** | **<0.0001** |
| | **PET date** | **0.0005** | **0.0001** | **4.2927** | **<0.0001** | | **PET date** | **0.0003** | **0.0001** | **2.0139** | **0.0441** |
| | head motion | 0.0021 | 0.0011 | 1.9366 | 0.0529 | | head motion | 0.0010 | 0.0013 | 0.7617 | 0.4463 |
| | CEN | 0.6711 | 0.2519 | 2.6644 | 0.0078 | | CEN | 0.2703 | 0.3141 | 0.8603 | 0.3897 |
| | **amyloid*CEN** | **0.0084** | **0.0028** | **3.0099** | **0.0026** | | **amyloid*CEN** | **0.0111** | **0.0034** | **3.2906** | **0.0010** |

Significant interactions or main effects in the absence of an interaction are bolded.

CEN - Central Executive Network, BGN - Basal Ganglia Network, DSST - digit symbol substitution test, eSPPB - expanded short physical performance battery

SE- standard error FDR - false discovery rate

Table S3. Adjusted models including age, brain volume measures, and head motion

| Outcome Variable | Independent Variables | Estimate | SE | T score | p-Value | Outcome Variable | Independent Variables | Estimate | SE | T score | p-Value |
|---|---|---|---|---|---|---|---|---|---|---|---|
| BGN | | | | | | | | | | | |
| Baseline DSST | amyloid | -0.0154 | 0.0460 | -0.3357 | 0.7371 | 30-month DSST | amyloid | -0.0924 | 0.0559 | -1.6547 | 0.0981 |
| | **age** | **0.1244** | **0.0389** | **3.1976** | **0.0014** | | **age** | **0.2658** | **0.0457** | **5.8095** | **<0.0001** |
| | gm_vol | 0.0017 | 0.0043 | 0.4059 | 0.6849 | | **gm_vol** | **-0.0111** | **0.0051** | **-2.1857** | **0.0289** |
| | **wm_vol** | **0.0144** | **0.0050** | **2.9128** | **0.0036** | | wm_vol | 0.0031 | 0.0061 | 0.5090 | 0.6108 |
| | **bilat_hippo** | **0.9118** | **0.2517** | **3.6229** | **0.0003** | | **bilat_hippo** | **2.2541** | **0.3033** | **7.4322** | **<0.0001** |
| | **bilat_thal** | **0.9534** | **0.2111** | **4.5174** | **<0.0001** | | bilat_thal | 0.3122 | 0.2545 | 1.2266 | 0.2201 |
| | bilat_pfc | 0.0237 | 0.0518 | 0.4579 | 0.6471 | | bilat_pfc | 0.0658 | 0.0617 | 1.0673 | 0.2859 |
| | **ICV** | **-0.0042** | **0.0014** | **-3.0108** | **0.0026** | | ICV | -0.0016 | 0.0017 | -0.9360 | 0.3493 |
| | **WMLV** | **0.0960** | **0.0316** | **3.0343** | **0.0024** | | WMLV | 0.0600 | 0.0377 | 1.5922 | 0.1114 |
| | head motion | 0.0034 | 0.0221 | 0.1533 | 0.8782 | | head motion | -0.0118 | 0.0260 | -0.4540 | 0.6499 |
| | **BGN** | **15.8255** | **5.1545** | **3.0702** | **0.0022** | | **BGN** | **16.1393** | **6.4289** | **2.5104** | **0.0121** |
| | amyloid*BGN | 0.0134 | 0.0600 | 0.2236 | 0.8231 | | amyloid*BGN | 0.1171 | 0.0726 | 1.6128 | 0.1069 |
| Baseline eSPPB | amyloid | -0.0049 | 0.0023 | -2.1483 | 0.0318 | 30-month eSPPB | amyloid | -0.0036 | 0.0028 | -1.2885 | 0.1977 |
| | **age** | **0.0203** | **0.0019** | **10.5957** | **<0.0001** | | **age** | **0.0192** | **0.0023** | **8.3331** | **<0.0001** |
| | gm_vol | 0.0002 | 0.0002 | 0.7452 | 0.4562 | | **gm_vol** | **0.0009** | **0.0003** | **3.3306** | **0.0009** |
| | wm_vol | 0.0000 | 0.0003 | -0.0187 | 0.9850 | | wm_vol | 0.0001 | 0.0003 | 0.2609 | 0.7942 |
| | bilat_hippo | 0.0076 | 0.0126 | 0.6002 | 0.5484 | | bilat_hippo | -0.0186 | 0.0154 | -1.2043 | 0.2286 |
| | bilat_thal | 0.0023 | 0.0106 | 0.2180 | 0.8274 | | bilat_thal | -0.0135 | 0.0128 | -1.0574 | 0.2904 |
| | bilat_pfc | -0.0045 | 0.0026 | -1.7076 | 0.0878 | | bilat_pfc | -0.0076 | 0.0031 | -2.4448 | 0.0146 |
| | ICV | 0.0000 | 0.0001 | 0.4446 | 0.6566 | | ICV | -0.0001 | 0.0001 | -1.0283 | 0.3039 |
| | **WMLV** | **0.0040** | **0.0016** | **2.5353** | **0.0113** | | **WMLV** | **0.0077** | **0.0021** | **3.5677** | **0.0004** |
| | **head motion** | **0.0025** | **0.0011** | **2.3410** | **0.0193** | | head motion | 0.0011 | 0.0013 | 0.8477 | 0.3967 |
| | **BGN** | **-0.0046** | **0.2605** | **-0.0176** | **0.9860** | | BGN | -0.0688 | 0.3215 | -0.2139 | 0.8306 |
| | **amyloid*BGN** | **0.0067** | **0.0030** | **2.2450** | **0.0248** | | amyloid*BGN | 0.0055 | 0.0036 | 1.5009 | 0.1335 |

Significant interactions or main effects in the absence of an interaction are bolded.

CEN - Central Executive Network, BGN - Basal Ganglia Network, DSST - digit symbol substitution test, eSPPB - expanded short physical performance battery

gm_vol - gray matter volume, wm_vol - white matter volume, bilat_hippo - bilateral hippocampal volume, bilat_thal - bilateral thalamus volume

bilat_pfc - bilateral prefrontal cortex volume, ICV - intracranial volume, WMLV - what matter lesion volume, SE- standard error FDR - false discovery rate

Table S3 cont. Adjusted models including age, brain volume measures, and head motion

| Outcome Variable | Independent Variables | Estimate | SE | T score | p-Value | Outcome Variable | Independent Variables | Estimate | SE | T score | p-Value |
|---|---|---|---|---|---|---|---|---|---|---|---|
| CEN | | | | | | | | | | | |
| Baseline DSST | amyloid | -0.0243 | 0.0399 | -0.6086 | 0.5428 | 30-month DSST | amyloid | -0.1168 | 0.0478 | -2.4449 | 0.0145 |
| | **age** | **0.1240** | **0.0387** | **3.2031** | **0.0014** | | **age** | **0.2656** | **0.0456** | **5.8294** | **<0.0001** |
| | gm_vol | 0.0002 | 0.0043 | 0.0441 | 0.9648 | | **gm_vol** | **-0.0126** | **0.0051** | **-2.4752** | **0.0134** |
| | **wm_vol** | **0.0135** | **0.0049** | **2.7221** | **0.0065** | | wm_vol | 0.0018 | 0.0061 | 0.3024 | 0.7623 |
| | **bilat_hippo** | **0.9432** | **0.2507** | **3.7618** | **0.0002** | | **bilat_hippo** | **2.2813** | **0.3024** | **7.5430** | **<0.0001** |
| | **bilat_thal** | **0.9225** | **0.2108** | **4.3773** | **<0.0001** | | bilat_thal | 0.2884 | 0.2543 | 1.1342 | 0.2568 |
| | bilat_pfc | 0.0282 | 0.0517 | 0.5447 | 0.5860 | | bilat_pfc | 0.0691 | 0.0615 | 1.1223 | 0.2618 |
| | **ICV** | **-0.0042** | **0.0014** | **-2.9951** | **0.0028** | | ICV | -0.0015 | 0.0017 | -0.9077 | 0.3641 |
| | **WMLV** | **0.0876** | **0.0316** | **2.7709** | **0.0056** | | WMLV | 0.0482 | 0.0377 | 1.2778 | 0.2014 |
| | head motion | -0.0066 | 0.0222 | -0.2999 | 0.7643 | | head motion | -0.0238 | 0.0262 | -0.9052 | 0.3654 |
| | **CEN** | **22.2809** | **5.1412** | **4.3338** | **<0.0001** | | CEN | 18.4032 | 6.2786 | 2.9311 | 0.0034 |
| | amyloid*CEN | 0.0273 | 0.0566 | 0.4820 | 0.6299 | | **amyloid*CEN** | **0.1626** | **0.0677** | **2.4027** | **0.0163** |
| Baseline eSPPB | amyloid | -0.0052 | 0.0020 | -2.6149 | 0.0090 | 30-month eSPPB | amyloid | -0.0067 | 0.0024 | -2.7900 | 0.0053 |
| | **age** | **0.0200** | **0.0019** | **10.5243** | **<0.0001** | | **age** | **0.0188** | **0.0023** | **8.2039** | **<0.0001** |
| | gm_vol | 0.0001 | 0.0002 | 0.3981 | 0.6906 | | **gm_vol** | **0.0008** | **0.0003** | **3.1776** | **0.0015** |
| | wm_vol | -0.0001 | 0.0003 | -0.2297 | 0.8183 | | wm_vol | 0.0000 | 0.0003 | 0.1448 | 0.8849 |
| | bilat_hippo | 0.0085 | 0.0126 | 0.6776 | 0.4981 | | bilat_hippo | -0.0189 | 0.0154 | -1.2279 | 0.2196 |
| | bilat_thal | 0.0011 | 0.0105 | 0.1058 | 0.9157 | | bilat_thal | -0.0138 | 0.0128 | -1.0783 | 0.2810 |
| | bilat_pfc | -0.0043 | 0.0026 | -1.6390 | 0.1013 | | bilat_pfc | -0.0075 | 0.0031 | -2.4178 | 0.0157 |
| | ICV | 0.0000 | 0.0001 | 0.4437 | 0.6573 | | ICV | -0.0001 | 0.0001 | -1.0296 | 0.3033 |
| | **WMLV** | **0.0036** | **0.0016** | **2.2796** | **0.0227** | | **WMLV** | **0.0071** | **0.0022** | **3.2881** | **0.0010** |
| | head motion | 0.0019 | 0.0011 | 1.7638 | 0.0779 | | head motion | 0.0006 | 0.0013 | 0.4687 | 0.6393 |
| | CEN | 0.7157 | 0.2533 | 2.8258 | 0.0047 | | CEN | 0.2259 | 0.3144 | 0.7184 | 0.4726 |
| | **amyloid*CEN** | **0.0076** | **0.0028** | **2.7379** | **0.0062** | | **amyloid*CEN** | **0.0103** | **0.0034** | **3.0547** | **0.0023** |

Significant interactions or main effects in the absence of an interaction are bolded.

CEN - Central Executive Network, BGN - Basal Ganglia Network, DSST - digit symbol substitution test, eSPPB - expanded short physical performance battery

gm_vol - gray matter volume, wm_vol - white matter volume, bilat_hippo - bilateral hippocampal volume, bilat_thal - bilateral thalamus volume

bilat_pfc - bilateral prefrontal cortex volume, ICV - intracranial volume, WMLV - what matter lesion volume, SE- standard error FDR - false discovery rate

Table S4. Sensitive analyses using the same 76 participants in all adjusted models

| Outcome Variable | Independent Variables | Estimate | SE | T score | p-Value | Outcome Variable | Independent Variables | Estimate | SE | T score | p-Value |
|---|---|---|---|---|---|---|---|---|---|---|---|
| BGN | | | | | | | | | | | |
| Baseline DSST | amyloid | -0.0385 | 0.0464 | -0.8302 | 0.4065 | 30-month DSST | amyloid | -0.1259 | 0.0565 | -2.2297 | 0.0258 |
| | **age** | **0.1520** | **0.0392** | **3.8789** | **0.0001** | | **age** | **0.2999** | **0.0461** | **6.4991** | **<0.0001** |
| | **PET date** | **0.0101** | **0.0023** | **4.3795** | **<0.0001** | | **PET date** | **0.0085** | **0.0029** | **2.9595** | **0.0031** |
| | head motion | 0.0097 | 0.0223 | 0.4376 | 0.6617 | | head motion | -0.0100 | 0.0263 | -0.3802 | 0.7038 |
| | **BGN** | **15.4589** | **5.2121** | **2.9660** | **0.0030** | | BGN | 15.1197 | 6.5100 | 2.3225 | 0.0203 |
| | amyloid*BGN | 0.0460 | 0.0605 | 0.7608 | 0.4468 | | **amyloid*BGN** | **0.1656** | **0.0733** | **2.2584** | **0.0240** |
| Baseline eSPPB | amyloid | -0.0050 | 0.0023 | -2.1803 | 0.0293 | 30-month eSPPB | amyloid | -0.0037 | 0.0028 | -1.3057 | 0.1918 |
| | **age** | **0.0205** | **0.0019** | **10.7966** | **<0.0001** | | **age** | **0.0194** | **0.0023** | **8.4766** | **<0.0001** |
| | **PET date** | **0.0006** | **0.0001** | **5.0652** | **<0.0001** | | **PET date** | **0.0005** | **0.0001** | **3.5748** | **0.0004** |
| | **head motion** | **0.0027** | **0.0011** | **2.5076** | **0.0122** | | head motion | 0.0016 | 0.0013 | 1.2074 | 0.2274 |
| | BGN | -0.0074 | 0.2594 | -0.0287 | 0.9771 | | BGN | -0.0585 | 0.3216 | -0.1818 | 0.8557 |
| | **amyloid*BGN** | **0.0068** | **0.0030** | **2.2898** | **0.0221** | | amyloid*BGN | 0.0055 | 0.0036 | 1.5255 | 0.1272 |
| CEN | | | | | | | | | | | |
| Baseline DSST | amyloid | -0.0424 | 0.0402 | -1.0549 | 0.2916 | 30-month DSST | amyloid | -0.1407 | 0.0483 | -2.9142 | 0.0036 |
| | **age** | **0.1507** | **0.0390** | **3.8662** | **0.0001** | | **age** | **0.2992** | **0.0459** | **6.5146** | **<0.0001** |
| | **PET date** | **0.0094** | **0.0023** | **4.0745** | **<0.0001** | | PET date | **0.0078** | **0.0029** | **2.7091** | **0.0068** |
| | head motion | -0.0026 | 0.0224 | -0.1150 | 0.9085 | | head motion | -0.0237 | 0.0265 | -0.8929 | 0.3720 |
| | **CEN** | **23.5162** | **5.1801** | **4.5397** | **<0.0001** | | CEN | 18.1023 | 6.3340 | 2.8580 | 0.0043 |
| | amyloid*CEN | 0.0557 | 0.0570 | 0.9778 | 0.3283 | | **amyloid*CEN** | **0.2018** | **0.0683** | **2.9543** | **0.0032** |
| Baseline eSPPB | amyloid | -0.0052 | 0.0020 | -2.6794 | 0.0074 | 30-month eSPPB | amyloid | -0.0070 | 0.0024 | -2.9262 | 0.0035 |
| | **age** | **0.0202** | **0.0019** | **10.7094** | **<0.0001** | | **age** | **0.0189** | **0.0023** | **8.3030** | **<0.0001** |
| | **PET date** | **0.0005** | **0.0001** | **4.8354** | **<0.0001** | | **PET date** | **0.0005** | **0.0001** | **3.4374** | **0.0006** |
| | head motion | 0.0020 | 0.0011 | 1.9017 | 0.0573 | | head motion | 0.0010 | 0.0013 | 0.7607 | 0.4469 |
| | CEN | 0.6777 | 0.2515 | 2.6952 | 0.0071 | | CEN | 0.2589 | 0.3133 | 0.8263 | 0.4087 |
| | **amyloid*CEN** | **0.0078** | **0.0028** | **2.8164** | **0.0049** | | **amyloid*CEN** | **0.0108** | **0.0034** | **3.1998** | **0.0014** |

Significant interactions or main effects in the absence of an interaction are bolded.

CEN - Central Executive Network, BGN - Basal Ganglia Network, DSST - digit symbol substitution test, eSPPB - expanded short physical performance battery

SE- standard error FDR - false discovery rate

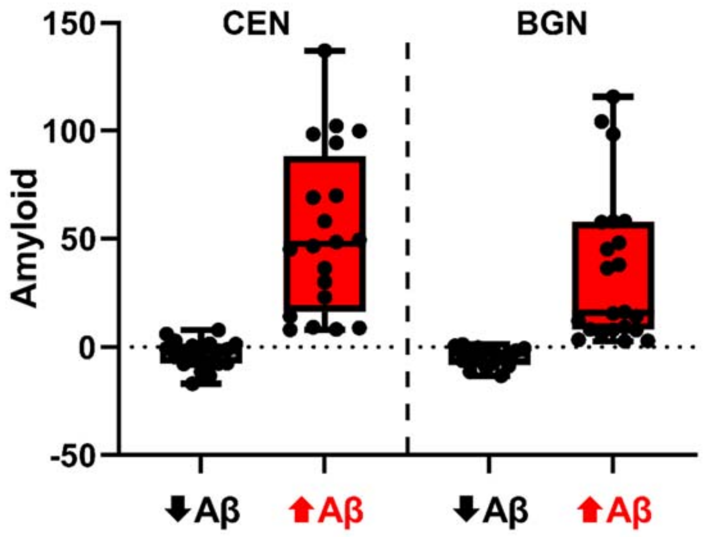


Figure S1: Box and whisker plots for amyloid levels (centiloids) for the high and low Aβ groups used to depict the interaction in Figure 4 of the main manuscript. The participants in each grouping are different based on the groupings for the CEN and BGN. The box depicts the boundaries of the upper and lower quartiles, and the line indicates the median.

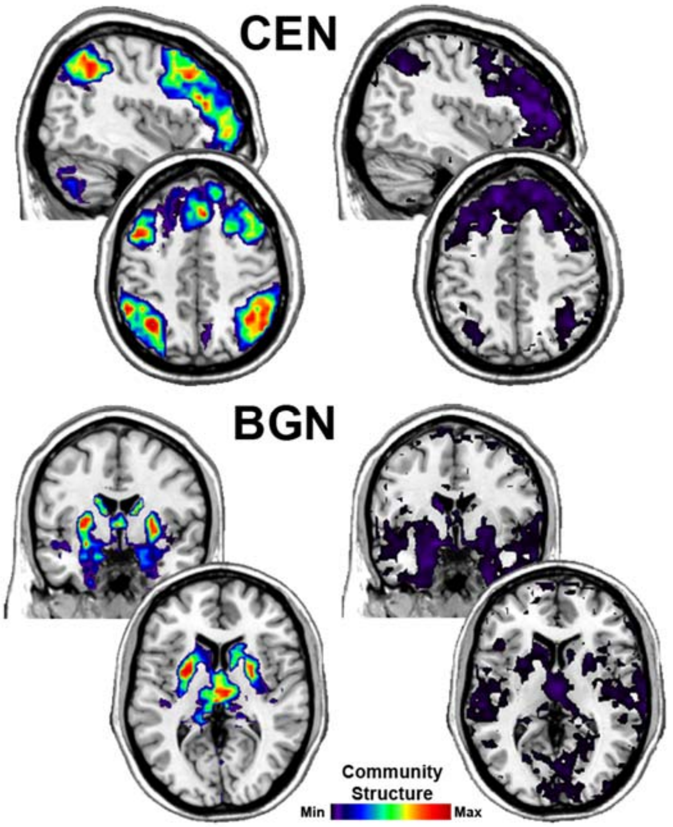


Figure S2: Average community structure images for the high (left) and low (right) network groups used to depict the interaction in Figure 4 of the main manuscript. The Montreal Neurological Institute (MIN) coordinates for the CEN are x = 42 for the upper sagittal slice and z = 46 for the lower axial slice. Coordinates for the BGN are y = -1 for the upper coronal slice and z = 6 for the lower axial slice. The calibration bar applies to all images but due to the vastly different scales, it was necessary to have a different range for each group to effectively visually depict the integrity of each network. The ranges are: CEN high (0.04-0.105) BGN high (0.02-0.05), low CEN (0.01-0.105), low BGN (0.035-0.06) all based on scaled inclusivity as described in the methods.
